# Population genetic simulation study of power in association testing across genetic architectures and study designs

**DOI:** 10.1101/632786

**Authors:** Dominic Ming Hay Tong, Ryan D. Hernandez

**Author notes:** Grant numbers: NIH 1R01HG007644.

## Abstract

While it is well established that genetics can be a major contributor to population variation of complex traits, the relative contributions of rare and common variants to phenotypic variation remains a matter of considerable debate. Here, we simulate rare variant association studies across different case/control panel sampling strategies, sequencing methods, and genetic architecture models based on evolutionary forces to determine the statistical performance of RVATs widely in use. We find that the highest statistical power of RVATs is achieved by sampling case/control individuals from the extremes of an underlying quantitative trait distribution. We also demonstrate that the use of genotyping arrays, in conjunction with imputation from a whole genome sequenced (WGS) reference panel, recovers the vast majority (90%) of the power that could be achieved by sequencing the case/control panel using current tools. Finally, we show that for dichotomous traits, the statistical performance of RVATs decreases as rare variants become more important in the trait architecture. Our results extend previous work to show that RVATs are insufficiently powered to make generalizable conclusions about the role of rare variants in dichotomous complex traits.

## Introduction

Genome-wide association studies (GWAS) have detected many common variants associated with hundreds of complex heritable phenotypes, but for many traits, much of that heritability remains unexplained. One proposed source of this so-called “missing heritability” are rare variants, which are hotly debated but have been implicated as a non-negligible source of genetic variance in prostate cancer (Mancuso et al., 2016), gene expression (Hernandez et al., 2017), height and BMI (Wainschtein et al., 2019). Unfortunately, power to detect rare variant associations is low in single-marker statistical tests at the genome-wide scale. Researchers have proposed many rare variant association tests (RVATs), statistical methods to pool rare variants within a putatively causal locus and test for association with the phenotype. These RVATs are broadly classified into three categories: burden tests (Liu & Leal, 2010), variance-component tests (Neale et al., 2011; Wu et al., 2011), and combined tests (Lee et al., 2012; Sun, Zheng, & Hsu, 2013). Though each test is published with its own validation simulations, these simulations are generally not comparable, and have their own flaws. (Moutsianas et al., 2015) systematically characterized the performance of commonly used gene-based rare variant association tests under a range of genetic architectures, sample sizes, variant effect sizes, and significance thresholds, and found that MiST, SKAT-O, and KBAC have the highest mean power across simulated data, but that these tests had overall low power even in the cases of loci with relatively large effect sizes.

It is well-known in the population genetics literature that population expansions and contractions (i.e. demography) can dramatically affect genome-wide patterns of genetic variation in a population (Auton et al., 2009; Bhaskar, Wang, & Song, 2015; Gravel et al., 2011; Uricchio, Zaitlen, Ye, Witte, & Hernandez, 2016), and that the action of natural selection can amplify or inhibit the frequency of functional alleles (Boyko et al., 2008; Eyre-Walker, Woolfit, & Phelps, 2006; Lohmueller et al., 2011). Together, these evolutionary forces shape the genetic architecture of complex traits (Lohmueller, 2014; Uricchio et al., 2016), and are critical components to understand in the pursuit of identifying the genetic basis for the bevy of human phenotypes understudy. Inferred demographic models of non-African human populations show a serial bottleneck model as populations migrated in waves across the globe, followed by explosive exponential growth since the dawn of agriculture. Moreover, studies of selection have found that most amino acid changes in proteins and changes in conserved non-coding loci are weakly deleterious (Boyko et al., 2008; Torgerson et al., 2009). Together, growth and selection has resulted in a preponderance of ultra-rare mutations (MAF<0.1%), which contribute a plurality of heritability in gene expression (Hernandez et al., 2017), BMI (Wainschtein et al., 2019), and possibly other traits. Accounting for demographic and selective effects on the frequency spectrum of causal variation is therefore crucial in characterizing the statistical power of RVATs. However, while previous evaluations of RVAT power have attempted to mimic the frequency spectrum of observed variants, they typically use phenotype models (or genetic architectures) that do not directly account for evolutionary forces like demography and natural selection and are often biologically unrealistic [e.g. effect sizes that are simple functions of the minor allele frequency (Wu et al., 2011)], limited to specific relative risks (Wray & Goddard, 2010), or lack pleiotropy (Moutsianas et al., 2015).

Another vital component of designing genetic association studies is the method of acquiring genetic data. Although the gold standard for capturing rare variation remains deep whole genome sequencing (WGS), the $1000 per genome cost still means performing WGS on any sizeable group of individuals remains prohibitively expensive for all but the largest consortia. Genotyping arrays make acquiring genetic data for a large number of individuals significantly less expensive, but lack coverage of rare variation. With larger WGS reference panels like the Haplotype Reference Consortium (HRC; McCarthy et al., 2016), large numbers of genotyped samples can be imputed to gain some insight into rare variation. With such large reference panels, imputation accuracy of genetic variation down to MAF≅0.1% is near perfect in European individuals (Quick et al., 2019). As more diverse reference panels become available [e.g. TOPMed (Taliun et al., 2019)], imputation in non-European and admixed populations will also improve, particularly for rare variants. Capturing these rare variants using genotyping arrays and imputation is more cost-effective and can lead to many more individuals in a study. However, imputation is limited by the variants that are carried by the individuals in the reference panel, and by the accuracy of the algorithm being used. Imputation accuracy falls off at lower minor allele frequencies (MAF), but the use of large WGS reference panels reduces the threshold of acceptable imputation quality (r^2^>0.3) to ~0.004-0.006% (Taliun et al., 2019) in European and African populations. Despite these limitations, imputation has been used to identify rare variant associations in acute macular degeneration (Helgason et al., 2013), lipid levels in type 2 diabetes patients (Marvel et al., 2017), systemic lupus erythematosus (Martinez-Bueno & Alarcón-Riquelme, 2019), among others. It is possible that additional rare variant association signals can be found in imputed data as imputation quality improves, but it is unclear what the statistical properties of RVATs in this setting are.

Here, we evaluate the statistical power of rare variant association tests in a simulation study under different genetic architectures, methods of acquiring genetic data, and methods of selecting individuals to be a part of the case-control cohort. We demonstrate how statistical power of RVATs is dependent on genetic architecture as well as sampling strategy for the case/control cohort. In particular, we find that sampling the extremes of a quantitative phenotype has the highest RVAT power, but power erodes quickly for all sampling strategies as the amount of genetic variance explained by rare variants increases.

## Materials and Methods

### Simulating genomic sequence data

We simulate neutral genetic sequence data under a coalescent model using msprime (Kelleher, Etheridge, & McVean, 2016) with a European and African demographic history (Tennessen et al., 2012). Under this demographic model, the European population experienced a series of bottlenecks as they moved out of Africa and into Europe. These bottlenecks were followed by super exponential growth in the European population and recent exponential growth in the African population, along with bi-directional migration. Using this neutral demographic model, we generate a 5Mb region with a mutation rate of 1e-8 and with genetic map arbitrarily chosen to mimic chr22:17000000-22000000 in hg19.

### Simulating genotype data

Some analyses are based on genotype array data. To simulate a genotyping array, we downsample the simulated neutral sequence data above to match the allele frequency spectrum and the average distance between variants of the Illumina OmniExpress2.5 genotyping chip, used in the GoT2D study (Fuchsberger et al., 2016).

### Simulating quantitative phenotypes

We transform our simulated neutral genetic data into quantitative phenotypes using a three-step procedure, following Uricchio et al (Uricchio et al., 2016). First, we simulate functional variants using the forward simulator SFS_CODE (Hernandez, 2008) under the same demographic model as above, but with purifying selection. Specifically, we generate 2000 independent loci of length 100kbp (for a total of 200Mb) with 100,000 individuals, where new mutations receive a fitness effect drawn from a gamma distribution [as inferred for non-synonymous sites (Boyko et al., 2008)]. This procedure generates a large table of functional variants, with corresponding derived allele counts and fitness effects.

The second step is to project the allele frequencies of our list of functional variants down to the desired sample size (using a binomial model), and transform fitness effects to phenotypic effect sizes using the Uricchio et al. model (Uricchio et al., 2016). This model parameterized the correlation between fitness effects and phenotypic effect sizes (through ρ) and the functional relationship between fitness effects and phenotypic effect sizes (through *τ* and *δ*). In particular, a causal variant with fitness effect *s* will have effect size *z*_*s*_ as follows:

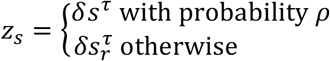

Under this model, with probability *ρ*, the effect size *z*_*s*_ of a site is a direct function of the site’s fitness effect (*s*), otherwise the effect size *z*_*s*_ is a function of a randomly sampled fitness effect (*s*_*r*_) drawn from the entire list of functional variants generated by the first step above. In this model, *δ* is +1 or −1 with equal probability to enable trait-increasing and trait-decreasing effects.

The third step for generating quantitative phenotypes is to identify the desired number of causal loci in our 5mb simulated sequence. For each variant within the causal loci, we sample a random variant from our list of functional variants generated in step two with the exact same allele frequency, and assign derived alleles at this causal site the effect size of the sampled functional variant. The quantitative phenotype of each individual (*Y*_*i*_, for the ith individual) is then generated under an additive model by summing the effect sizes of all causal alleles that they carry:

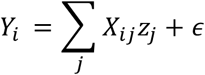

Where *Z*_*j*_ is the effect size of causal variant *j*, *X*_*ij*_ is the number of causal alleles carried by individual *i* at site *j*, and *ε* is a Normal random variable with mean 0 and variance 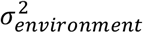 (which ensures the desired level of heritability of the trait). See Table 1 for the specific values of ρ, and heritability that are evaluated in this study. In contrast to previous work with this phenotypic model (Uricchio et al., 2016), we will focus on dichotomous traits, and describe our sampling strategy for such traits below.

**Table 1.**
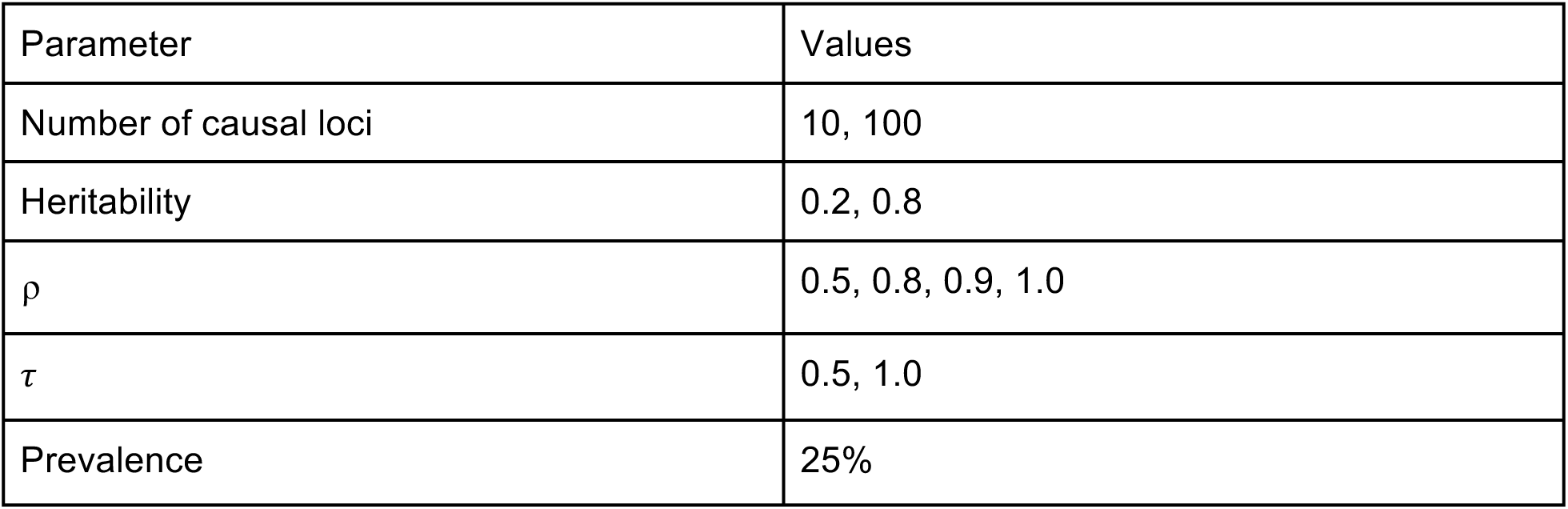
Genetic architectures examined in this study

| Parameter | Values |
| --- | --- |
| Number of causal loci | 10, 100 |
| Heritability | 0.2, 0.8 |
| $\rho$ | 0.5, 0.8, 0.9, 1.0 |
| $\tau$ | 0.5, 1.0 |
| Prevalence | 25% |

**Table 2.**
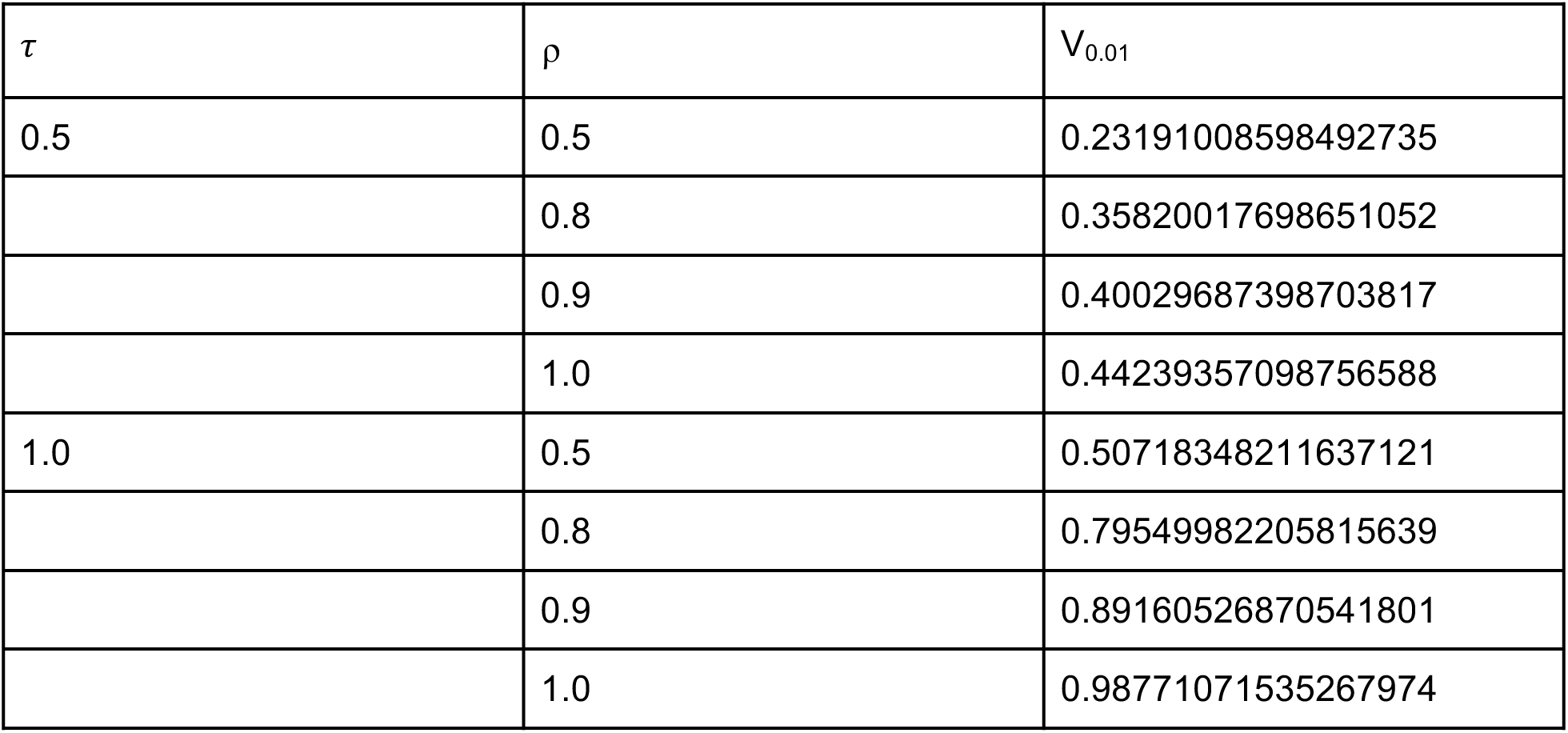
Genetic architecture parameters under the Uricchio model and the genetic variance explained by variants under MAF=1%

| $\tau$ | $\rho$ | $V_{0.01}$ |
| --- | --- | --- |
| 0.5 | 0.5 | 0.23191008598492735 |
|  | 0.8 | 0.35820017698651052 |
|  | 0.9 | 0.40029687398703817 |
|  | 1.0 | 0.44239357098756588 |
| 1.0 | 0.5 | 0.50718348211637121 |
|  | 0.8 | 0.79549982205815639 |
|  | 0.9 | 0.89160526870541801 |
|  | 1.0 | 0.98771071535267974 |

**Table 3.** Study design parameters in this study

| Parameter | Value |
| --- | --- |
| Sampling strategy | Random, 50/50, extremes |
| Number of case/control individuals | 5000, 10000 |
| Number of reference panel individuals for imputing | 10000, 20000 |

### Selecting sampling strategies for association tests

The quantitative phenotypes can be dichotomized to simulate three different sampling strategies: random, 50/50, and extremes. In the extreme sampling strategy, we sample the desired number of individuals from the top and bottom of the quantitative phenotype distribution. For the random and 50/50 sampling strategy, we first define the individuals with quantitative phenotypes in the top P% to be our population of cases (where P represents the prevalence of our trait of interest), and the remaining individuals to be our population of controls. We then sample cases and controls from their respective populations. For the random sampling strategy, we sample cases in proportion to the prevalence of the trait, while for the 50/50 sampling strategy we sample equal numbers of cases and controls. The random sampling strategy is used as a worst case scenario to establish the worst possible power under that sampling strategy.

### Imputing genotyped data

In some analyses, we evaluate the effectiveness of genotype imputation. Such analyses require two data sets: the phenotype sample (e.g. case/control or continuous phenotype), and an imputation reference panel. The dichotomized phenotype individuals are generated as above, with their genetic data down-sampled to mimic a genotype array platform. We then sample an additional set of individuals from the total population to form the imputation panel. The down-sampled genotype data is then pre-phased using SHAPEIT2 (Delaneau, Marchini, & Zagury, 2012) and imputed using IMPUTE4 (Bycroft et al., 2017).

### Running tests of association on simulated data

We ran rare variant association tests (RVATs) using the rvtests software (Zhan, Hu, Li, Abecasis, & Liu, 2016). We focus on SKAT (Wu et al., 2011), SKAT-O (Lee et al., 2012), and KBAC (Liu & Leal, 2010), which were found to be most powerful in detecting disease-associated variation in a previous study (Moutsianas et al., 2015). We applied each RVAT to non-overlapping analysis blocks of 10kbp across the simulated region, and computed power and false-positive rates for each test as the proportion of simulations with p-values below 2.5e-6. We ran logistic regression on each variant above MAF=1% to determine associations with the phenotype using PLINK. The detection threshold was set at 5e-8. To compare GWAS to RVAT power, we evaluate if there is a variant under the GWAS p-value threshold within the 10kb analysis block. If there is such a variant, we deem the GWAS to have found that analysis block to be causal for comparisons with RVAT.

### Calculating cumulative genetic variance

We follow (Uricchio et al., 2016) in calculating *V*_*x*_, the genetic variance due to variants at or below allele frequency *x*, which is given by:

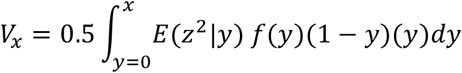

Where *f*(*y*)is the site frequency spectra (SFS), i.e. the proportion of sampled alleles at frequency *y*, and *E*(*z*^2^|*y*) is the mean-squared effect size of variants at frequency *y*. We pool 20 simulations of 300kbp in 50k African individuals using msprime to obtain an accurate measure of the SFS and the expected effect size of variants at frequency *x*. To normalize across genetic architectures, we divide by *V*_1_, which is the total additive genetic variance. The *V*_0.01_/*V*_1_ values (denoted as just *V*_0.01_ below) are used to denote the degree to which rare variants (variants with MAF ≤ 1%) matter under a particular pair of parameters under the Uricchio genetic architectures.

### Dataset and software availability

All scripts and datasets generated in this study, along with the results of single variant and gene-based association tests, are available on the website github.com/dmctong/rv_imp.

## Results

### Rare variants explain a majority of heritability only under restrictive scenarios

To determine whether there is genetic variance explained by rare variants, we calculated the expected genetic variance analytically under different (ρ,τ) combinations of the Uricchio model being studied here (see Methods and Table 1). In Figure 1, we show that the proportion of genetic variance explained as a function of MAF. We focus on the genetic variance explained by variants with MAF < 1% (V_0.01_), which varies dramatically between 99% when ρ=1, τ=1, to less than 1% when ρ=0, τ=0.5. We note that when τ=1 and ρ≠0, rare variants constitute a substantial fraction of the genetic variance (V_0.01_ > 40%), and the majority of the rare variant contribution is explained by singletons in this simulated sample of 50,000 individuals. In contrast, when τ=0.5 and ρ≠0, V_0.01_ ranges from ~20%-60% but singletons are expected to make a more subtle contribution to the genetic architecture of the trait.

**Figure 1.**
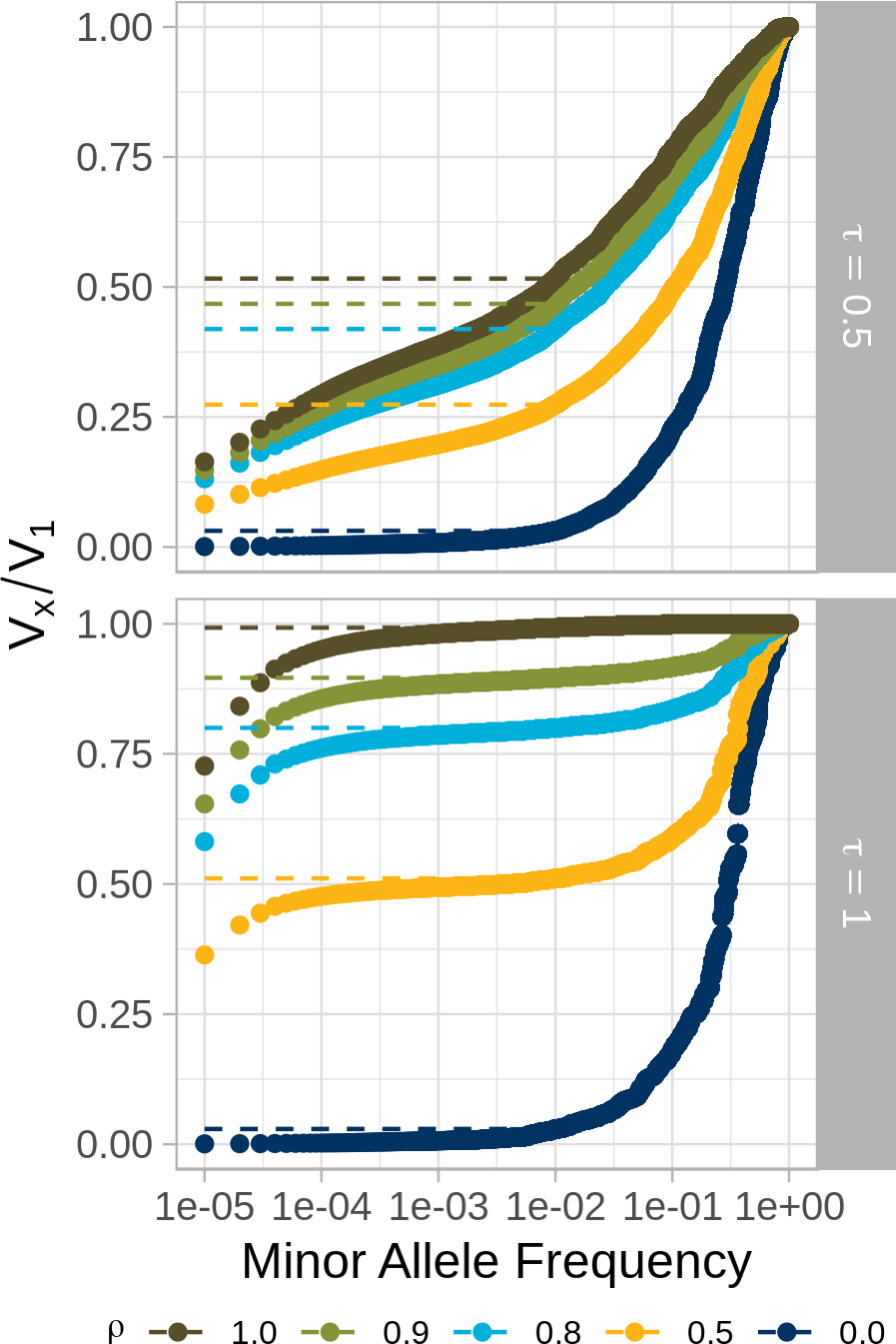
The cumulative proportion of the genetic variance explained by variants under minor allele frequency x (V_x_/V_1_) for a sample of 5000 individuals drawn from an African population demographic model under different values of ρ and τ in the Uricchio model. Top: τ = 0.5; bottom: τ = 1. Dotted lines indicate the proportion of genetic variance explained by alleles under 1% MAF (referred to as *V*_0.01_).

**Figure 2.**
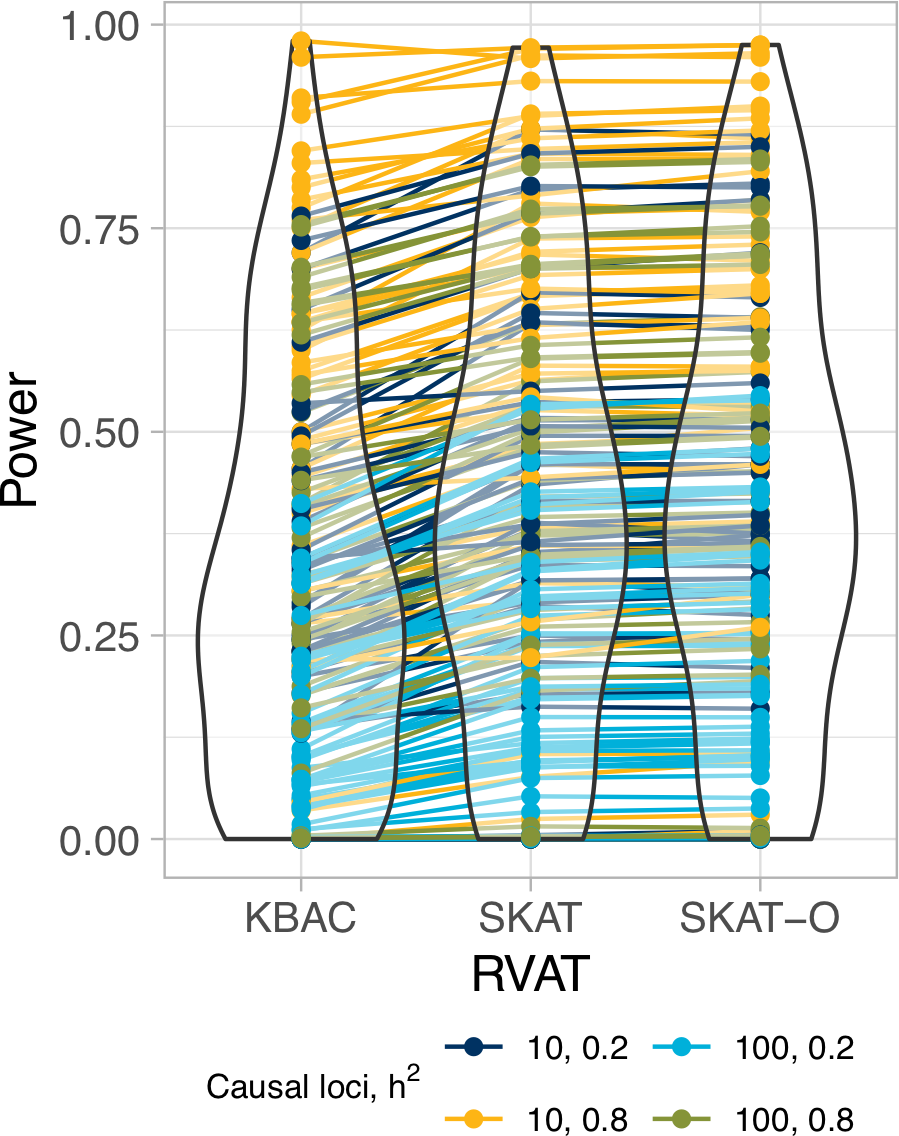
A global overview of the statistical power of a burden test (KBAC), a variance-component test (SKAT), and a combined test (SKAT-O) for all parameters shown in Table 1 using 50,000 African individuals simulated under an Out-of-Africa demographic model. Each point represents a genetic architecture tested with 10 independent simulations under the RVAT indicated; lines connect the same simulated parameters across RVATs to show that, generally speaking, the rank of statistical power is preserved across RVATs.

### Statistical power varies dramatically across different study designs, genetic architectures, and polygenicity, but not across RVATs

Figure 1 shows that rare variants can contribute substantial heritability to a trait under certain genetic architectures. Now we ask if we can detect the loci that harbor the causal rare variants using existing RVATs. To quantify the effects of genetic architecture and study design on the statistical power of RVATs, we focus on KBAC, SKAT, and SKAT-O, which represent each of the three major categories of RVATs and have been shown to be among the most powerful (Moutsianas et al., 2015). For the 5Mb region we simulated (see Methods), we raster over parameters in genetic architecture (heritability, number of causal loci, and the relationship between selection and phenotypic effect sizes; Table 1) and in study design (sequencing vs genotype imputation and selection of individuals in the case/control vs extreme phenotype panels). In Figure 2, we show the global overview of statistical power across all simulations. We find that the statistical power of all three RVATs is similar regardless of simulated parameters, but tend to be highest with SKAT and SKAT-O [p_MWU_(SKAT, SKAT-O)=0.795; p_MWU_(KBAC, SKAT-O)=0.002304]. As expected, power is higher when the causal signal is more concentrated (e.g. when heritability is high or the effect sizes are large due to few causal loci). Given the correspondence among tests, we will focus on SKAT-O in further analyses.

**Figure 3.**
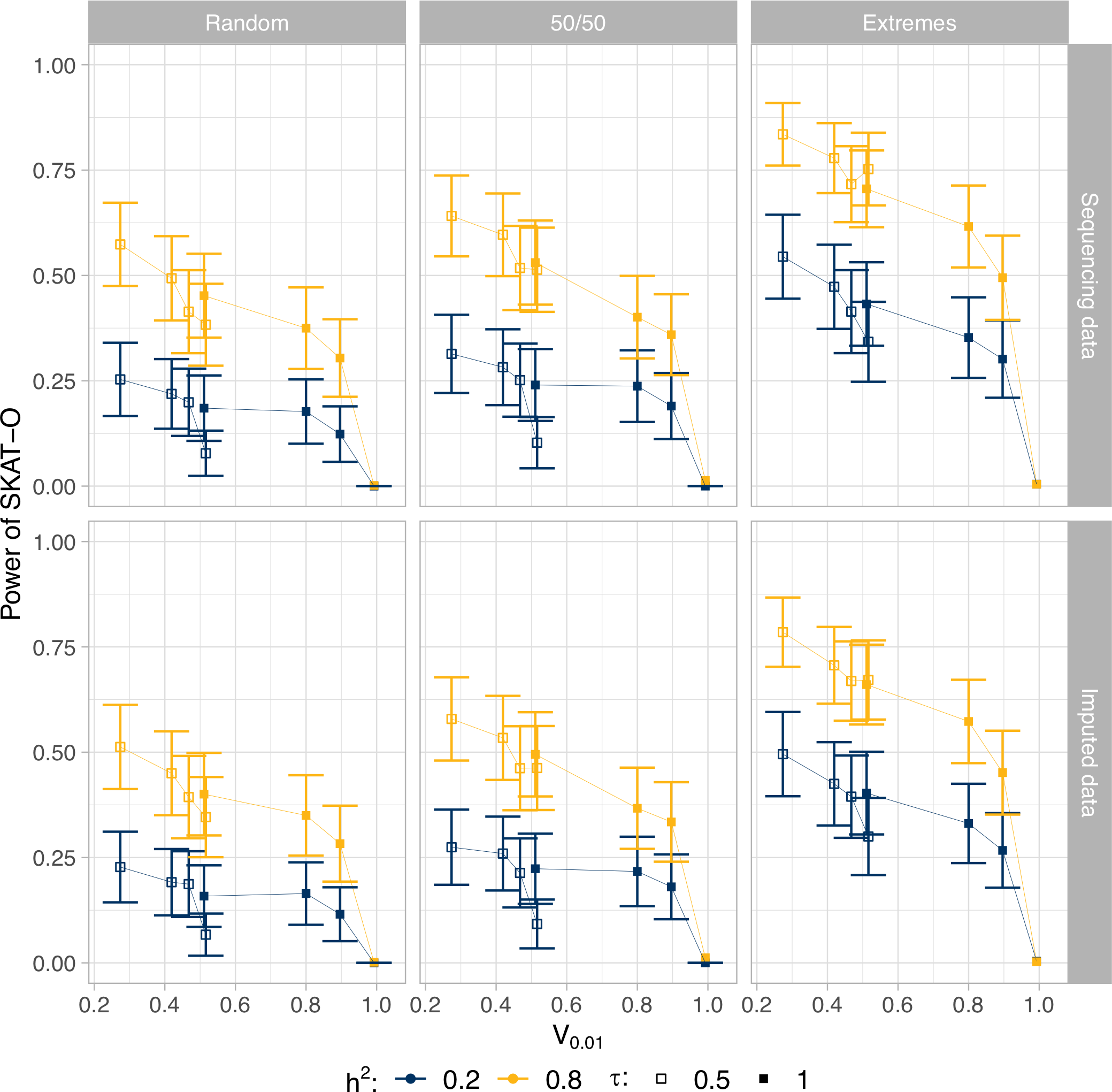
The statistical power of SKAT-O across different sampling strategies (columns) and across different sequencing methods (rows), as a function of the proportion of genetic variance explained by that genetic architecture at MAF=1%. Each point represents 20 independent simulations of 100 causal loci of 10kb each across a 5Mb simulated region for a given genetic architecture for a 50,000 individual African population.

### As rare variants explain more genetic variance of the trait, SKAT-O power decreases

We then ask how SKAT-O power changes as a function of genetic architecture. In Figure 3, we show that as V_0.01_ increases (i.e., as rare variants explain increasing amounts of the genetic variance of the trait), the power of SKAT-0 decreases. This pattern holds across all sampling strategies and for different levels of polygenicity (Figure S3). These results show patterns that will repeat in future sections: the extremes study design demonstrates the best overall power, followed by 50/50 and then random. Further, a more concentrated signal (higher heritability and/or lower number of causal loci, see Supplemental Figures) improves power. We found that as the functional form relating effect size to selection coefficient changes from τ=0.5 to τ=1, power increases slightly again, suggesting that V_0.01_ may be an overly simplistic characterization of the genetic architecture. Finally, applying SKAT-O to imputed data (bottom facet) reproduces all of the patterns we see when RVATs are applied to sequencing data (top facet), albeit with slightly worse power.

### Using extreme cases and controls as a sampling strategy improves statistical power of SKAT-O

The number of individuals sequenced as part of a study is a key design parameter of that study. To understand how increasing the number of individuals improves the statistical power of SKAT-O, we simulate across genetic architectures and study designs to find the increase in power per individual using SKAT-O from 2,500 to 20,000 individuals (Figure 4). In the extremes study design, where half of the individuals in the panel are selected from the extreme cases and half of the individuals are selected from the extreme controls (from a total population of 50,000 individuals), we find that mean power gain is zero. Increasing the number of individuals in this design means more individuals are drawn from closer to the mean of the distribution, so power is already maximized with a smaller sample of 2,500 individuals (and may actually decrease under some scenarios). In the random and 50/50 study designs, increasing the size of the case/control panel increases the number of relevant individuals, and so mean power gain is approximately 2e-5 per individual added.

**Figure 4.**
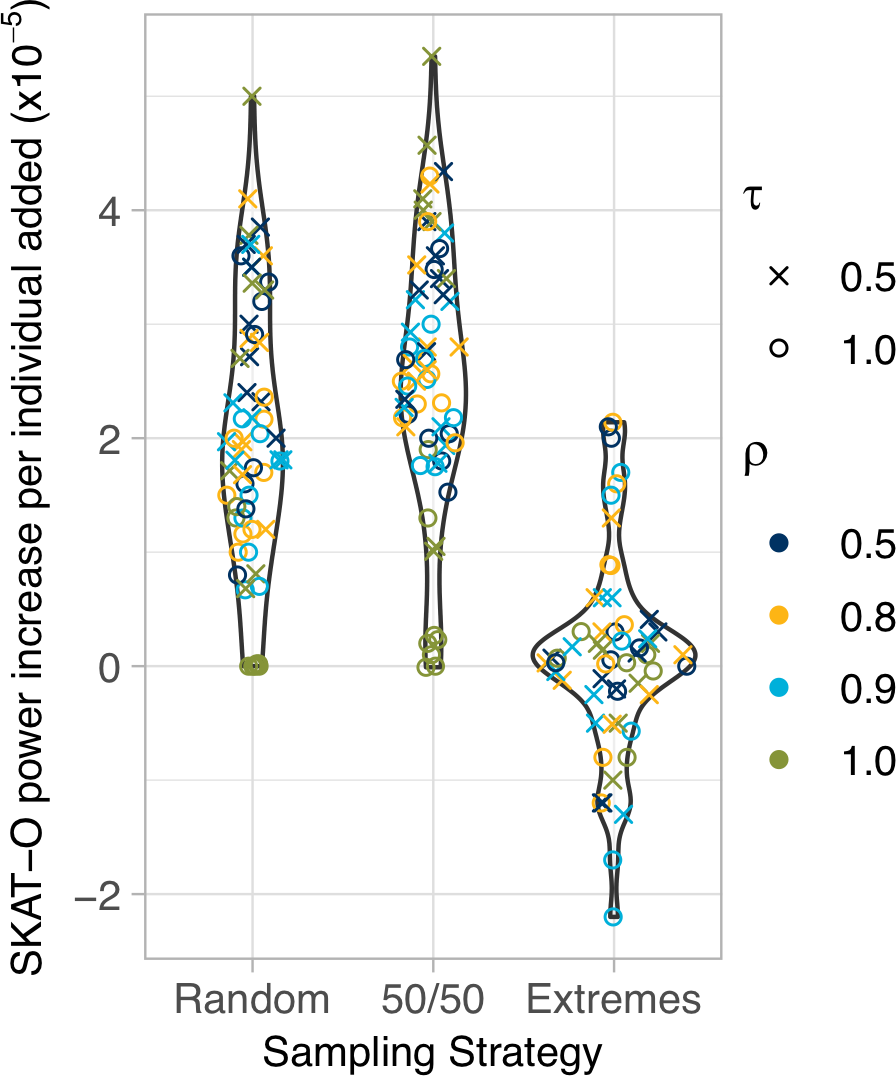
Increase in SKAT-O power as a function of sample size. SKAT-O power increases when increasing the sample size in non-extreme sampling strategies. Each point represents the slope from increasing the number of individuals in the case/control panel under a simulated genetic architecture.

This increase is highly dependent on the genetic architecture underlying the trait of interest.

### RVATs perform nearly as well on imputed data as they do on sequence data

Most genetic association studies have started with genotyping arrays to collect genomic data, followed by imputation against WGS reference panels to maximize discovery potential with single variant analyses. As WGS cost falls, more studies will conduct large-scale WGS, but here we ask if there is a potential opportunity to discover rare variant associations with imputed data. In Figure 5, we compare the mean power of SKAT-O when applied to genotyped-then-imputed samples to the mean power of SKAT-O applied to sequencing data from the same samples. We find that the decrease in power is minimal. Indeed, we find a robust linear relationship between RVAT power with sequencing vs imputed data, suggesting that for all scenarios evaluated here, imputation loses 10% power, on average, compared to sequencing data.

**Figure 5.**
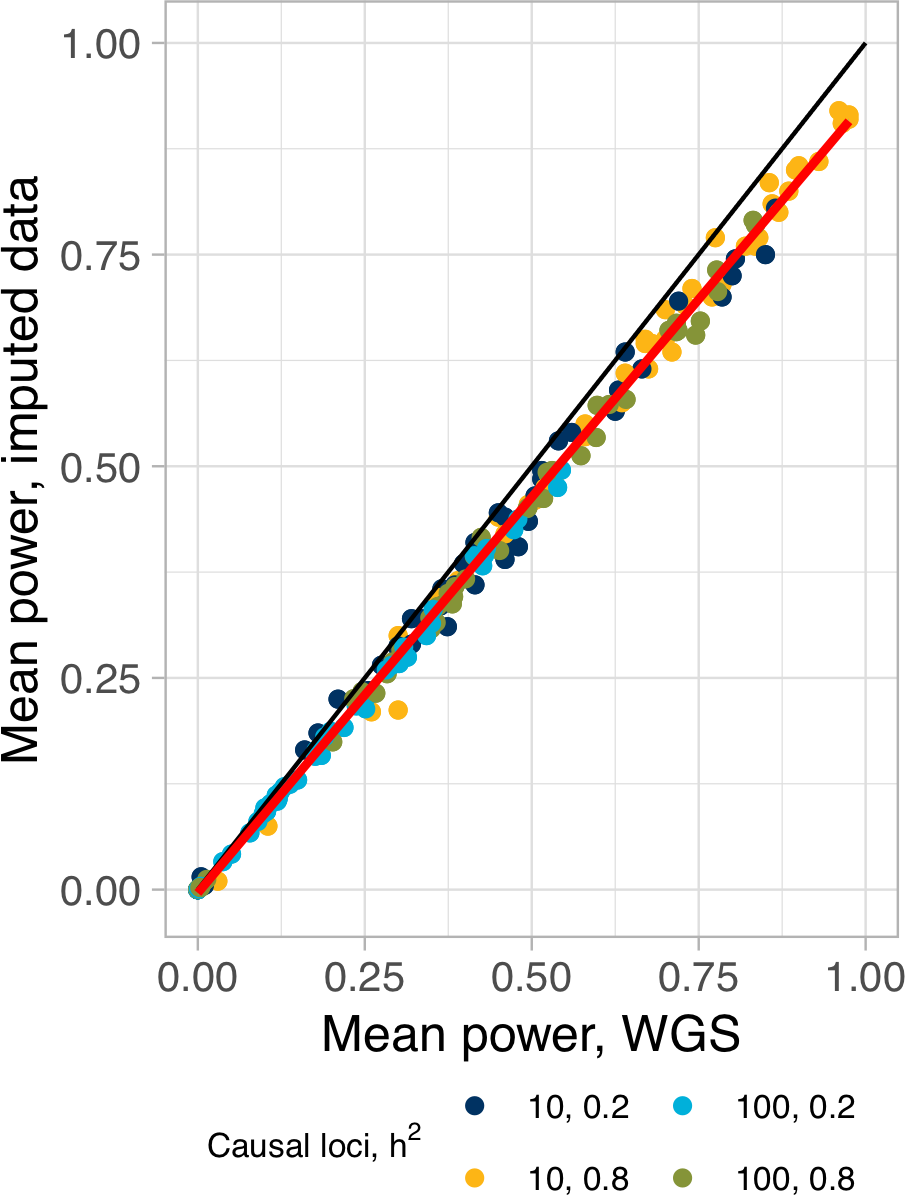
The mean power of SKAT-O across different genetic architectures using imputed data, compared to using sequence data. Each point represents a different simulated genetic architecture where we vary the number of causal bins (10 or 100), heritability (0.2 or 0.8), sampling strategy, (ρ, τ) for the underlying phenotype distribution, and the number of simulated case/control individuals in the study.

### RVATs under a GWAS peak

The general process of discovering genetic associations typically begins with genotyping and imputing a sample of individuals, followed by GWAS. The (typically unknown) genetic architecture of the trait determines the likelihood that a common variant will be detected with GWAS, and whether a rare variant association signal should be expected. Rastering over parameters of our phenotype model, a genome-wide significant single marker association (GWAS) was identified at 44.4% of causal loci. Figure 6 shows the power of SKAT-O using sequencing or imputed data conditional on seeing (circles) or not seeing (x’s) a statistically significant GWAS hit at a causal locus. We find that under all phenotype model parameters and sampling strategies evaluated, when a GWAS hit is identified, SKAT-O has at least 70% power to detect a rare variant signal with sequence data (and slightly less power with imputed data). If no GWAS peak is identified, there is considerably less power to identify a rare variant signal (and power further erodes as the genetic variance explained by rare variants increases).

We then mimic the process of first doing locus discovery on a sample of imputed individuals followed by sequencing for different sampling strategies. In Figure 7, we show that sequencing data has at least 75% power to replicate causal loci identified with imputed data (regardless of the genetic architecture and case-control sampling strategy). However, when no association is found with imputed data, power to identify causal loci with sequencing data is highly dependent on the case-control sampling strategy, and the overall heritability and genetic architecture of the trait (with power generally decreasing as V_0.01_ increases).

**Figure 6.**
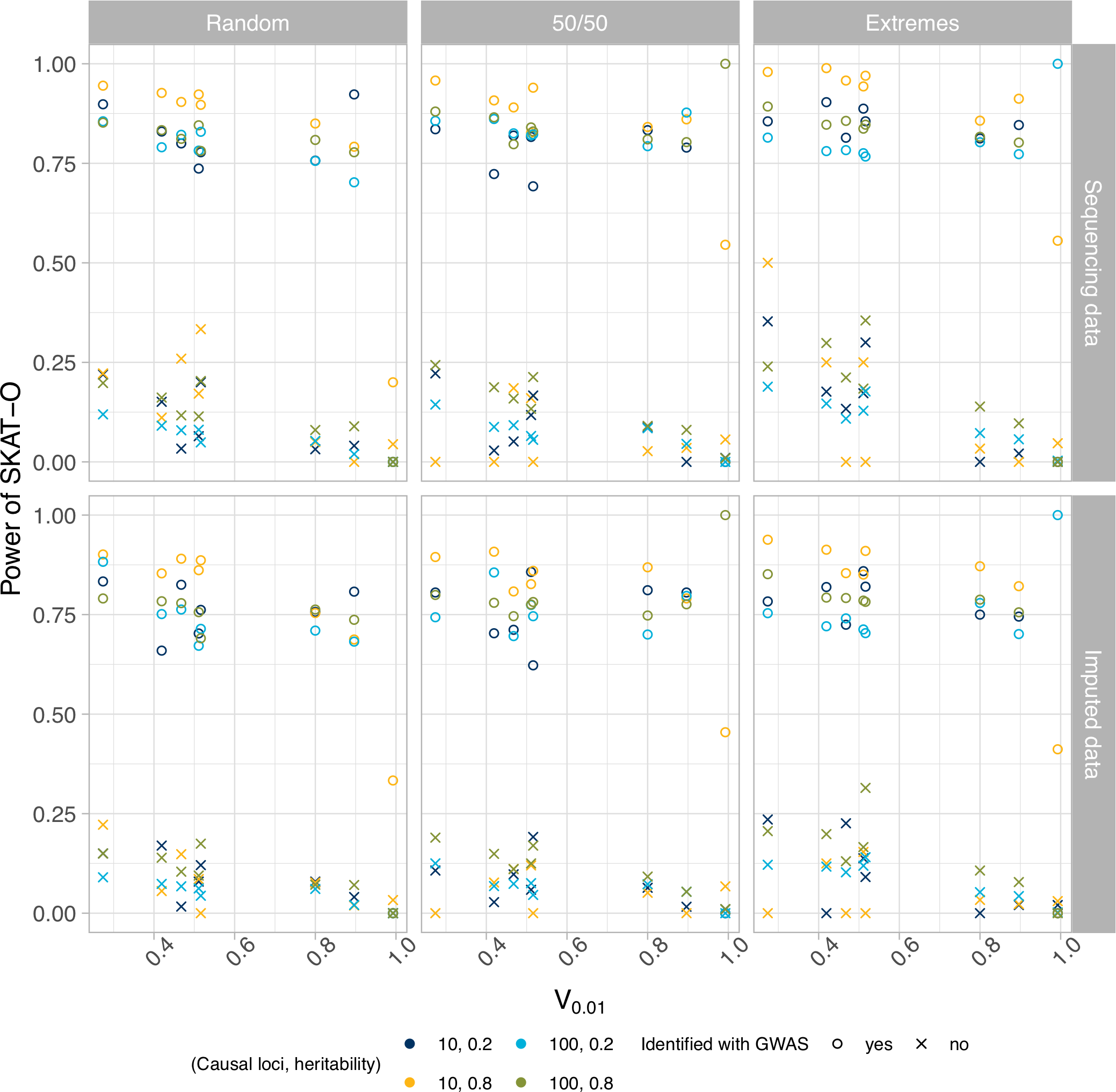
The statistical power of GWAS given the results of SKAT-O, across different sequencing methods (rows) and across different sampling strategies (columns), as a function of the cumulative genetic variance explained by variants under 1% minor allele frequency. The shape shows the prediction of SKAT-O; the colours show the underlying number of causal loci and heritability of the trait.

**Figure 7.**
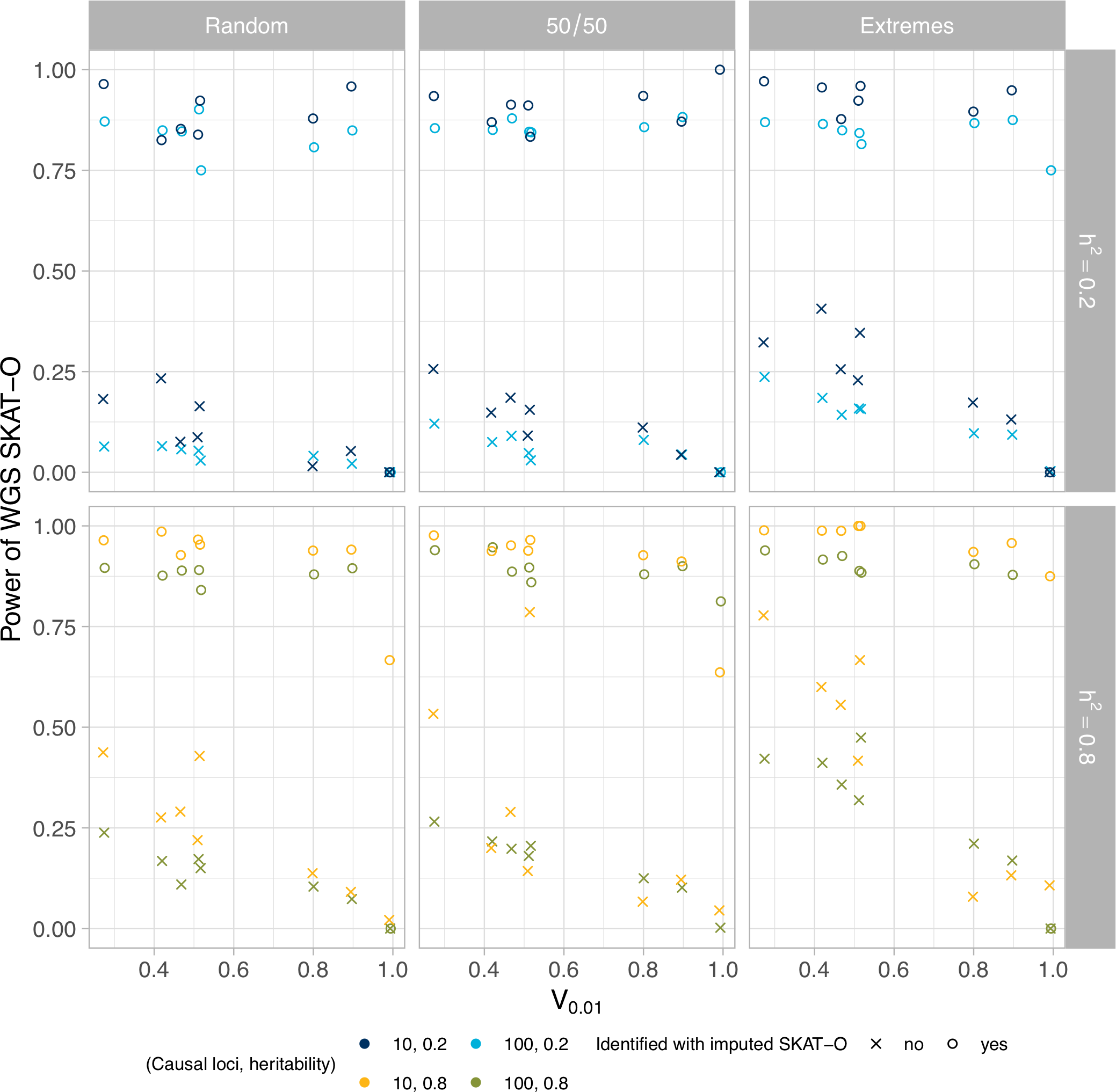
SKAT-O power using sequencing data, given the results of SKAT-O applied to imputed data. The shape indicates whether SKAT-O applied to imputed data correctly identified the causal locus (circles) or missed it (x). The colours show the underlying causal number of loci and heritability of the trait.

### Window of discovery around causal loci

In Figure 8, we plot the probability of SKAT-O detecting an association signal as a function of the distance from a causal locus. To benchmark the width of this discovery window, we use the full-width half-maximum statistic, which is the distance at which the probability of a significant association crosses below 50% of its maximum value (i.e. falls below 50% of the power estimated at the causal locus). Consistent with previous results, the full-width half-maximum is largest when there is a large amount of heritability concentrated in few causal loci and under the extremes study design. The larger points in Figure 8 represent this window of discovery, which is, on average, 34.3kb (sd 18.4kb) in the random study design, 42.8kb (sd 19.3kb) in the 50/50 design, and 64.3kb (sd 34.2kb) in the extremes design.

**Figure 8.**
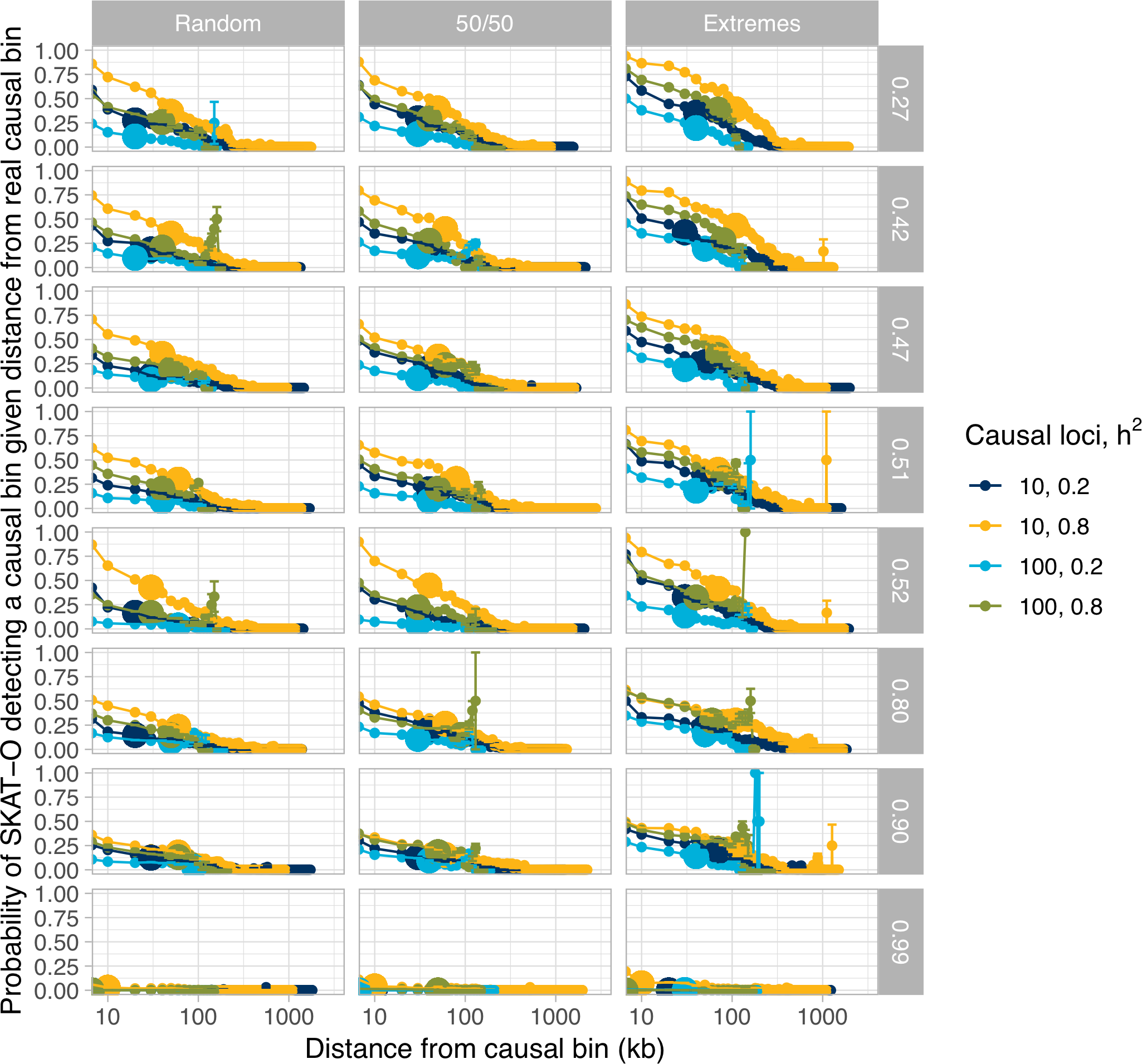
The window of discovery around causal loci, shown as the fraction of simulations that result in a statistically significant RVAT p-value as a function of distance from the nearest causal locus. Different sampling strategies are shown in columns, and V_0.01_ thresholds are shown in rows. Error bars are binomial standard errors of the mean. Bigger points represent full-width half-maximum points.

## Discussion

Genome-wide association studies (GWAS) so far have produced thousands of SNP associations for hundreds of traits (Witte, 2010). However, in these GWAS, the associated SNPs do not recapitulate the estimated heritability of the trait, leading to the problem of “missing heritability”. Though there are many proposed sources of this missing heritability, one popular hypothesis is that this missing heritability resides in rare variants. This has led to the development of rare variant association tests and massive investment in large whole genome sequencing studies. With these tests and this data becoming more and more prevalent, we look at how to optimize the design of a rare variant association study to maximize power.

It is clear that RVATs can be very powerful for detecting associations under simple genetic architectures [like when the effect size is proportional to log10(MAF) as proposed by (Wu et al., 2011)]. Such phenotype models do not take into account evolutionary forces like natural selection and demography, and it is well appreciated that genetic architectures are sensitive to these non-equilibrium evolutionary forces (Gazave, Chang, Clark, & Keinan, 2013; Simons, Turchin, Pritchard, & Sella, 2014). Uricchio et al presented a phenotype model that accounts for selection and pleiotropy and showed that existing RVATs struggle at realistic variance explained in genes across different human demographic histories (Uricchio et al., 2016). The Uricchio model captures modularity through the parameter ρ and the relationship between selection and effect size through τ, which enables a thorough exploration of different genetic architectures a trait could have (Figure 1).

We showed analytically that there is a significant amount of genetic variance explained in rare variants across different (ρ, τ) parameterizations under the Uricchio model (Figure 1), particularly when τ is equal to 1. These results are not surprising, as it has been shown that a substantial amount of heritability derives from rare variants in real traits like gene expression (Hernandez et al., 2017), height and BMI (Wainschtein et al., 2019). Taken together, the significant amount of heritability explained by rare variants under different parameterizations of the Uricchio model shows that RVATs have the potential to associate much of the causal variation underlying a complex trait. However, this model has only been studied in the context of continuous traits. We extend this model to study dichotomous traits (with case/control and extreme phenotype sampling strategies).

Many existing rare variant association tests were thoroughly characterized by (Moutsianas et al., 2015). We chose the most powerful representatives of the three classes of RVATs to use in our study: a variance-component test (SKAT), a burden method (KBAC), and a combined method (SKAT-O). Across all genetic architectures and study designs, we found that SKAT-O is the best performer, so we used SKAT-O in all further analyses on RVAT power in a case/control association study.

To run a case/control association study, the first step is to determine which individuals to select for your study, and how to acquire their genetic data. We simulated three different sampling strategies: randomly sampling cases and controls proportional to the trait prevalence; sampling half of your study size from cases and half from your controls; and sampling individuals from the extreme tails of a quantitative distribution [or a proxy underlying the trait such as bronchodilator response (Spear et al., 2018), for example]. Our results show that choosing from the tails of an underlying quantitative distribution produces the best power. This means for any case/control association study, spending some time to find the extreme tails of an underlying quantitative distribution for a trait will likely produce the best possible RVAT power.

We considered two ways of acquiring genetic data: using a genotyping array followed by imputation against a large reference panel, and direct sequencing of your study sample. Although a $1,000 whole genome is now possible, over the sample sizes required for an effective rare variant association study, the cost is prohibitive except for the largest consortia. Using genotyping arrays then imputing is still much less expensive than WGS (Quick et al., 2019), which could enable more than 5x more genotyped samples than WGS samples.

Applying SKAT-O to imputed data is expected to have lower power for several reasons. First, imputation accuracy decreases as MAF decreases (Howie, Marchini, Stephens, & Chakravarti, 2011; Quick et al., 2019), meaning fewer rare variants will be accurately imputed and correctly identified in the study sample. Second, imputation accuracy is highest when the study sample population and the reference panel population match, and this is not guaranteed to be the case, particularly when the study sample is from a minority population or an admixed population. Third, a majority of rare variants carried by the imputed samples are unlikely to be carried by the reference panel.

Comparing SKAT-O power across genetic architectures and study designs, we show that genotyping then imputing is about 90% as powerful as WGS using the same number of individuals. This implies that using genotyping then imputing with a larger sample size could produce as much if not more power than a smaller WGS sample. For most current rare variant association studies, our results suggest that using genotyping then imputing is the best way to start. We also looked at the increase in SKAT-O power using WGS after running a genotyping and imputation study; there is a boost in SKAT-O power when using WGS data following imputed data, but the trade-off between cost and power is something to be considered on an individual study basis.

The next step in characterizing RVAT power is to consider the genetic architecture of the trait of interest. Though complex trait architectures are not thoroughly understood, we used the Uricchio model to simulate different architectures and label these architectures using the amount of cumulative genetic variance explained by all variants under 1% minor allele frequency (V_0.01_). We show that SKAT-O power decreases as V_0.01_ increases, meaning SKAT-O performance is worst when rare alleles make the largest contributions to trait variance. Although counterintuitive, as one would expect RVATs are best tuned for the scenarios where rare variants matter most in the genetic architecture, our result mirrors the findings of Uricchio et al, 2016. One explanation is that as V_0.01_ goes up, the proportion of V_0.01_ due to singletons and other ultra-rare alleles increases as well, and statistically associating these ultra-rare alleles is difficult in the RVAT frameworks we evaluated here. We also note that the explosive exponential growth of the Tennessen demographic model used to simulate genetic data leads to an excess of ultra-rare alleles compared to the neutral expectation, such that both cases and controls harbor many ultra-rare variants (thereby confounding RVAT power).

With the decrease in power as rare variants mattered more, we wondered whether nearby regions in rare variant-dominated architectures would provide additional information. We looked at how the probability of SKAT-O detecting a causal region decreases as a function of distance from a causal region. The results suggest that in an unbiased window-based approach to scanning the genome with SKAT-O, positive hits that are not in causal regions may be useful in helping identify true causal regions, although again only in genetic architectures where rare variants do not contribute the majority of genetic variance. Interestingly, the power ranking of study designs is inverse of the ranking of precision, meaning that with higher power comes a larger window of discovery.

We also looked at the statistical properties of a common analytical path from GWAS to RVATs, and from imputed data to sequence data. We found that GWAS and SKAT-O are generally concordant, with causal regions identified by GWAS being identified by SKAT-O, while a smaller proportion (~15%) of causal regions are identified by SKAT-O and not by GWAS. We see little downside in testing for causal regions using SKAT-O following GWAS, with the ability to pick up additional causal regions on the same data. We caution that this effect declines significantly as rare variants explain more of the genetic variance.

Finally, the number of loci contributing to a trait (or its polygenicity) may be another important component of the trait’s architecture. It is not surprising that we found that for a fixed heritability of a trait, RVAT discovery power is higher when there are fewer true causal loci (as effect sizes are concentrated into fewer variants). However, it is possible that the polygenicity of a trait could be constraining the possible range of genetic architectures.

This study has a few limitations. It is based on simulated data that matches inferred human evolutionary history (including selection, and demographic history) but these models and simulations are incomplete representations of nature. We do not explore the effects of gene size, mutation rate, haplotype length, or degree of linkage disequilibrium between causal regions. We do not consider the differences between coding and non-coding regions, which have different selection coefficient distributions and potentially different contributions to the genetic architectures for a trait. Future work should consider a phenotype model where the function of a region is taken into account, as ENCODE (The ENCODE Project Consortium, 2012) and other consortia are rapidly adding more dimensions to genomic data. One major shortcoming is that we analyze only African and European populations in this study. With significant growth in admixed populations already happening - the US Census in 2014-15 predicts that the US will be a “majority-minority” country by 2050 (*Projections of the Size and Composition of the U.S. Population: 2014 to 2060*, 2014), meaning significant growth in African-American and Latino populations - it will be important to study association testing power in admixed populations. We also believe that incorporating functional annotations, evolutionary forces, and admixture into rare variant association tests would significantly improve statistical power.

## Acknowledgments

We thank Jeffrey D. Wall, Ian Holmes, and Raul Torres for stimulating discussions. This work was partially supported by the National Institutes of Health (grant 1R01HG007644 to R.D.H.). D.M.H.T. was partially supported by a National Sciences and Engineering Research Council of Canada Postgraduate Scholarships-Doctoral PGSD3-471524-2015. We thank Kevin Hartman for providing a useful script. We declare no conflicts of interest.

## References

Auton, A., Bryc, K., Boyko, A. R., Lohmueller, K. E., Novembre, J., Reynolds, A., … Bustamante, C. D. (2009). Global distribution of genomic diversity underscores rich complex history of continental human populations. Genome Research, 19(5), 795–803. https://doi.org/10.1101/gr.088898.108

Bhaskar, A., Wang, Y. X. R., & Song, Y. S. (2015). Efficient inference of population size histories and locus-specific mutation rates from large-sample genomic variation data. Genome Research, 25(2), 268–279. https://doi.org/10.1101/gr.178756.114

Boyko, A. R., Williamson, S. H., Indap, A. R., Degenhardt, J. D., Hernandez, R. D., Lohmueller, K. E., … Bustamante, C. D. (2008). Assessing the Evolutionary Impact of Amino Acid Mutations in the Human Genome. PLoS Genetics, 4(5), e1000083. https://doi.org/10.1371/journal.pgen.1000083

Bycroft, C., Freeman, C., Petkova, D., Band, G., Elliott, L. T., Sharp, K., … O’Connell, J. (2017). Genome-wide genetic data on~ 500,000 UK Biobank participants. BioRxiv, 166298.

Delaneau, O., Marchini, J., & Zagury, J.-F. (2012). A linear complexity phasing method for thousands of genomes. Nature Methods, 9(2), 179–181. https://doi.org/10.1038/nmeth.1785

Eyre-Walker, A., Woolfit, M., & Phelps, T. (2006). The Distribution of Fitness Effects of New Deleterious Amino Acid Mutations in Humans. Genetics, 173(2), 891–900. https://doi.org/10.1534/genetics.106.057570

Fuchsberger, C., Flannick, J., Teslovich, T. M., Mahajan, A., Agarwala, V., Gaulton, K. J., … McCarthy, M. I. (2016). The genetic architecture of type 2 diabetes. Nature, 536(7614), 41–47. https://doi.org/10.1038/nature18642

Gazave, E., Chang, D., Clark, A. G., & Keinan, A. (2013). Population Growth Inflates the Per-Individual Number of Deleterious Mutations and Reduces Their Mean Effect. Genetics, 195(3), 969–978. https://doi.org/10.1534/genetics.113.153973

Gravel, S., Henn, B. M., Gutenkunst, R. N., Indap, A. R., Marth, G. T., Clark, A. G., … McVean, G. A. (2011). Demographic history and rare allele sharing among human populations. Proceedings of the National Academy of Sciences, 108(29), 11983–11988. https://doi.org/10.1073/pnas.1019276108

Helgason, H., Sulem, P., Duvvari, M. R., Luo, H., Thorleifsson, G., Stefansson, H., … Stefansson, K. (2013). A rare nonsynonymous sequence variant in C3 is associated with high risk of age-related macular degeneration. Nature Genetics, 45(11), 1371–1374. https://doi.org/10.1038/ng.2740

Hernandez, R. D. (2008). A flexible forward simulator for populations subject to selection and demography. Bioinformatics, 24(23), 2786–2787. https://doi.org/10.1093/bioinformatics/btn522

Hernandez, R. D., Uricchio, L. H., Hartman, K., Ye, J., Dahl, A., & Zaitlen, N. (2017). Singleton Variants Dominate the Genetic Architecture of Human Gene Expression. BioRxiv, 219238.

Howie, B., Marchini, J., Stephens, M., & Chakravarti, A. (2011). Genotype Imputation with Thousands of Genomes. G3&#58; Genes|Genomes|Genetics, 1(6), 457–470. https://doi.org/10.1534/g3.111.001198

Kelleher, J., Etheridge, A. M., & McVean, G. (2016). Efficient coalescent simulation and genealogical analysis for large sample sizes. PLoS Computational Biology, 12(5), e1004842.

Lee, S., Emond, M. J., Bamshad, M. J., Barnes, K. C., Rieder, M. J., Nickerson, D. A., … Lin, X. (2012). Optimal Unified Approach for Rare-Variant Association Testing with Application to Small-Sample Case-Control Whole-Exome Sequencing Studies. The American Journal of Human Genetics, 91(2), 224–237. https://doi.org/10.1016Zj.ajhg.2012.06.007

Liu, D. J., & Leal, S. M. (2010). A Novel Adaptive Method for the Analysis of Next-Generation Sequencing Data to Detect Complex Trait Associations with Rare Variants Due to Gene Main Effects and Interactions. PLoS Genetics, 6(10), e1001156. https://doi.org/10.1371/journal.pgen.1001156

Lohmueller, K. E. (2014). The Impact of Population Demography and Selection on the Genetic Architecture of Complex Traits. PLoS Genetics, 10(5), e1004379. https://doi.org/10.1371/journal.pgen.1004379

Lohmueller, K. E., Albrechtsen, A., Li, Y., Kim, S. Y., Korneliussen, T., Vinckenbosch, N., … Nielsen, R. (2011). Natural Selection Affects Multiple Aspects of Genetic Variation at Putatively Neutral Sites across the Human Genome. PLoS Genetics, 7(10), e1002326. https://doi.org/10.1371/journal.pgen.1002326

Martínez-Bueno, M., & Alarcón-Riquelme, M. E. (2019). Exploring Impact of Rare Variation in Systemic Lupus Erythematosus by a Genome Wide Imputation Approach. Frontiers in Immunology, 10, 258. https://doi.org/10.3389/fimmu.2019.00258

Marvel, S. W., Rotroff, D. M., Wagner, M. J., Buse, J. B., Havener, T. M., McLeod, H. L., … The ACCORD/ACCORDion Investigators. (2017). Common and rare genetic markers of lipid variation in subjects with type 2 diabetes from the ACCORD clinical trial. PeerJ, 5, e3187. https://doi.org/10.7717/peerj.3187

McCarthy, S., Das, S., Kretzschmar, W., Delaneau, O., Wood, A. R., Teumer, A., … Marchini, J. (2016). A reference panel of 64,976 haplotypes for genotype imputation. Nature Genetics, 48(10), 1279–1283. https://doi.org/10.1038/ng.3643

Moutsianas, L., Agarwala, V., Fuchsberger, C., Flannick, J., Rivas, M. A., Gaulton, K. J., … McCarthy, M. I. (2015). The Power of Gene-Based Rare Variant Methods to Detect Disease-Associated Variation and Test Hypotheses About Complex Disease. PLOS Genetics, 11(4), e1005165. https://doi.org/10.1371/journal.pgen.1005165

Neale, B. M., Rivas, M. A., Voight, B. F., Altshuler, D., Devlin, B., Orho-Melander, M., … Daly, M. J. (2011). Testing for an Unusual Distribution of Rare Variants. PLoS Genetics, 7(3), e1001322. https://doi.org/10.1371/journal.pgen.1001322

Projections of the Size and Composition of the U.S. Population: 2014 to 2060. (2014). 13.

Quick, C., Anugu, P., Musani, S., Weiss, S. T., Burchard, E. G., White, M. J., … Fuchsberger, C. (2019). Sequencing and Imputation in GWAS: Cost-Effective Strategies to Increase Power and Genomic Coverage Across Diverse Populations [Preprint]. https://doi.org/10.1101/548321

Simons, Y. B., Turchin, M. C., Pritchard, J. K., & Sella, G. (2014). The deleterious mutation load is insensitive to recent population history. Nature Genetics, 46(3), 220–224. https://doi.org/10.1038/ng.2896

Spear, M. L., Hu, D., Pino-Yanes, M., Huntsman, S., Eng, C., Levin, A. M., … Burchard, E. G. (2018). A genome-wide association and admixture mapping study of bronchodilator drug response in African Americans with asthma. The Pharmacogenomics Journal. https://doi.org/10.1038/s41397-018-0042-4

Sun, J., Zheng, Y., & Hsu, L. (2013). A Unified Mixed-Effects Model for Rare-Variant Association in Sequencing Studies. Genetic Epidemiology, 37(4), 334–344. https://doi.org/10.1002/gepi.21717

Taliun, D., Harris, D. N., Kessler, M. D., Carlson, J., Szpiech, Z. A., Torres, R., … Abecasis, G. R. (2019). Sequencing of 53,831 diverse genomes from the NHLBI TOPMed Program [Preprint]. https://doi.org/10.1101/563866

Tennessen, J. A., Bigham, A. W., O’Connor, T. D., Fu, W., Kenny, E. E., Gravel, S., … on behalf of the NHLBI Exome Sequencing Project. (2012). Evolution and Functional Impact of Rare Coding Variation from Deep Sequencing of Human Exomes. Science, 337(6090), 64–69. https://doi.org/10.1126/science.1219240

The ENCODE Project Consortium. (2012). An integrated encyclopedia of DNA elements in the human genome. Nature, 489(7414), 57–74. https://doi.org/10.1038/nature11247

Mancuso, N., Rohland, N., Rand, K. A., Tandon, A., Allen, A., … Reich, D. (2016). The contribution of rare variation to prostate cancer heritability. Nature Genetics, 48(1), 30–35. https://doi.org/10.1038/ng.3446

Torgerson, D. G., Boyko, A. R., Hernandez, R. D., Indap, A., Hu, X., White, T. J., … Clark, A. G. (2009). Evolutionary Processes Acting on Candidate cis-Regulatory Regions in Humans Inferred from Patterns of Polymorphism and Divergence. PLoS Genetics, 5(8), e1000592. https://doi.org/10.1371/journal.pgen.1000592

Uricchio, L. H., Torres, R., Witte, J. S., & Hernandez, R. D. (2015). Population Genetic Simulations of Complex Phenotypes with Implications for Rare Variant Association Tests. Genetic Epidemiology, 39(1), 35–44. https://doi.org/10.1002/gepi.21866

Uricchio, L. H., Zaitlen, N. A., Ye, C. J., Witte, J. S., & Hernandez, R. D. (2016). Selection and explosive growth alter genetic architecture and hamper the detection of causal rare variants. Genome Research, 26(7), 863–873. https://doi.org/10.1101/gr.202440.115

Wainschtein, P., Jain, D. P., Yengo, L., Zheng, Z., TOPMed Anthropometry Working Group, Trans-Omics for Precision Medicine Consortium, Cupples, L. A., … Visscher, P. M. (2019). Recovery of trait heritability from whole genome sequence data [Preprint]. https://doi.org/10.1101/588020

Witte, J. S. (2010). Genome-Wide Association Studies and Beyond. Annual Review of Public Health, 31(1), 9–20. https://doi.org/10.1146/annurev.publhealth.012809.103723

Wray, N. R., & Goddard, M. E. (2010). Multi-locus models of genetic risk of disease. Genome Medicine, 2(2), 10. https://doi.org/10.1186/gm131

Wu, M. C., Lee, S., Cai, T., Li, Y., Boehnke, M., & Lin, X. (2011). Rare-Variant Association Testing for Sequencing Data with the Sequence Kernel Association Test. The American Journal of Human Genetics, 89(1), 82–93. https://doi.org/10.1016/j.ajhg.2012.06.007

Zhan, X., Hu, Y., Li, B., Abecasis, G. R., & Liu, D. J. (2016). RVTESTS: an efficient and comprehensive tool for rare variant association analysis using sequence data: Table 1. Bioinformatics, 32(9), 1423–1426. https://doi.org/10.1093/bioinformatics/btw079

