## Supplemental Figures for "Population genetic simulation study of power in association testing across genetic architectures and study designs"

Dominic Ming Hay Tong[1], Ryan D. Hernandez[1,2,3]
[1]University of California, Berkeley – University of California, San Francisco Graduate Program in Bioengineering
[2]Department of Bioengineering and Therapeutic Sciences, University of California, San Francisco
[3]Department of Human Genetics, McGill University, Montreal, QC, Canada

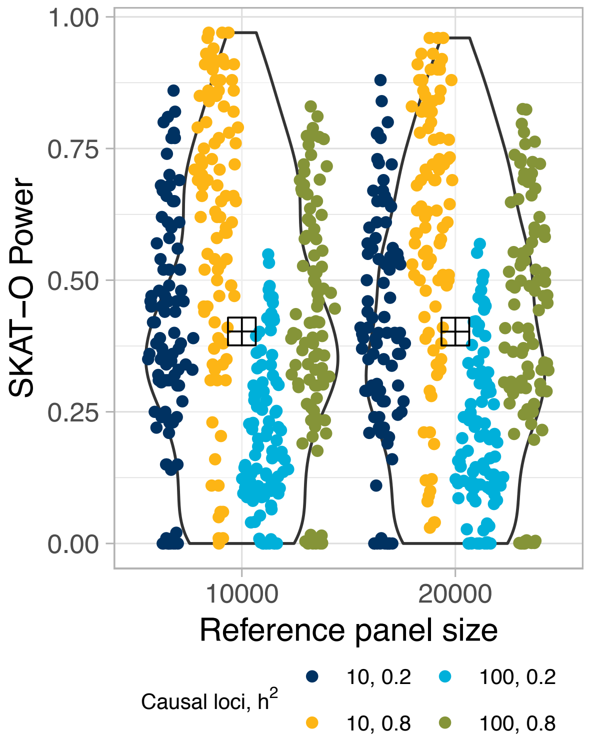


Figure S1. The statistical power of SKAT-O under an imputed study design does not change when the reference panel size is increased. Each point represents 10 independent simulations of a genetic architecture. The larger square point in the middle of each distribution indicates the mean SKAT-O power under each reference panel size.

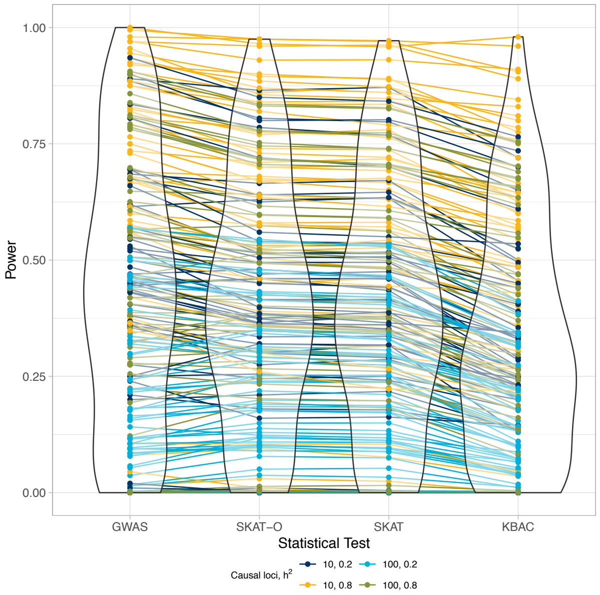


Figure S2.  A global overview of the statistical power of a burden test (KBAC), a variance-component test (SKAT), a combined test (SKAT-O), and a single-variant logistic regression test (GWAS) for all parameters shown in Table 1 using 50,000 African individuals simulated under an Out-of-Africa demographic model. Each point represents a genetic architecture tested with 10 independent simulations under the RVAT indicated; lines connect the same simulated parameters across RVATs to show that, generally speaking, the rank of statistical power is preserved across RVATs.

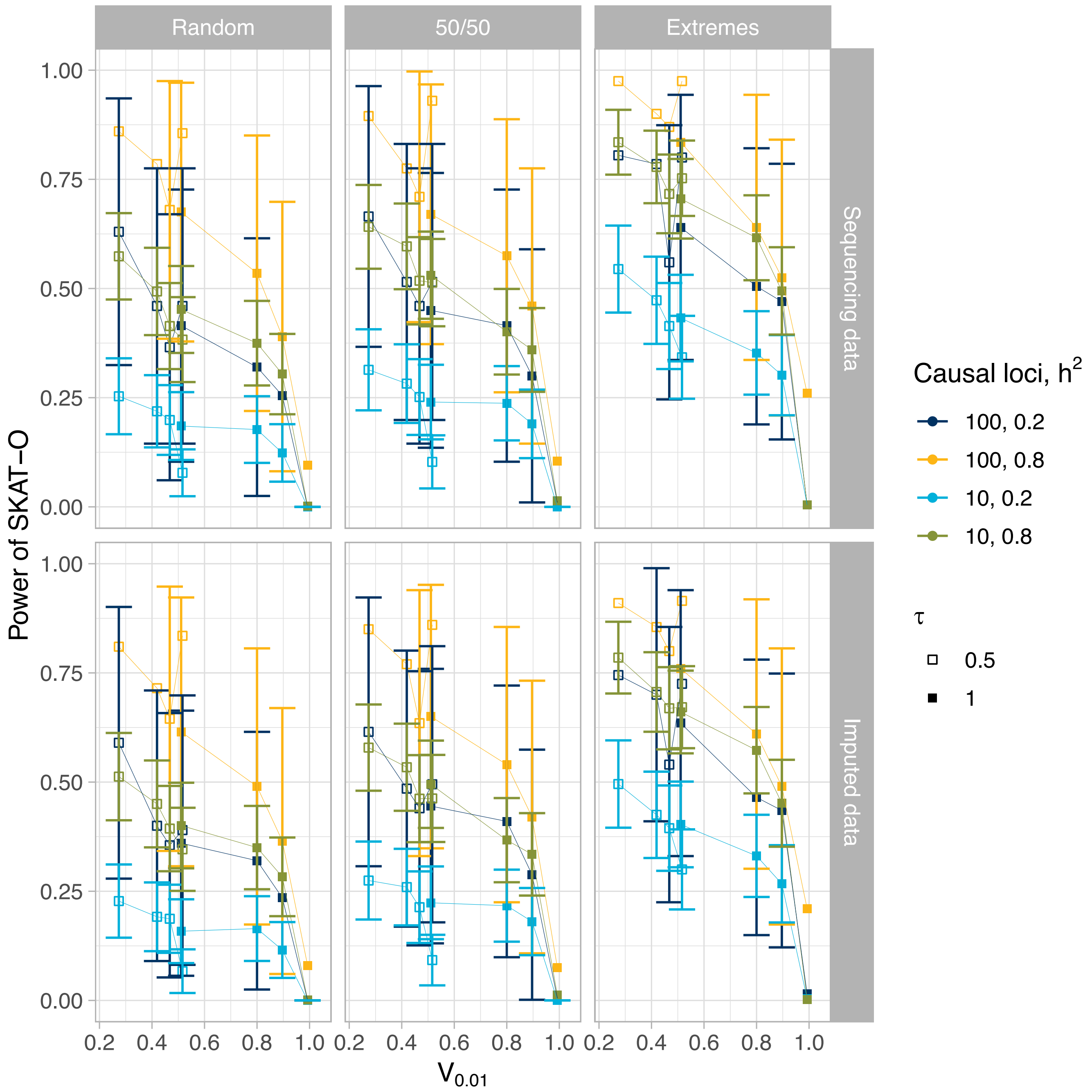


Figure S3. The statistical power of SKAT-O across different sampling strategies (columns) and across different sequencing methods (rows), as a function of the proportion of genetic variance explained by that genetic architecture at MAF=1%. Each point represents 20 independent simulations of 10 or 100 causal loci of 10kb each across a 5Mb simulated region for a given genetic architecture for a 50,000 individual African population.

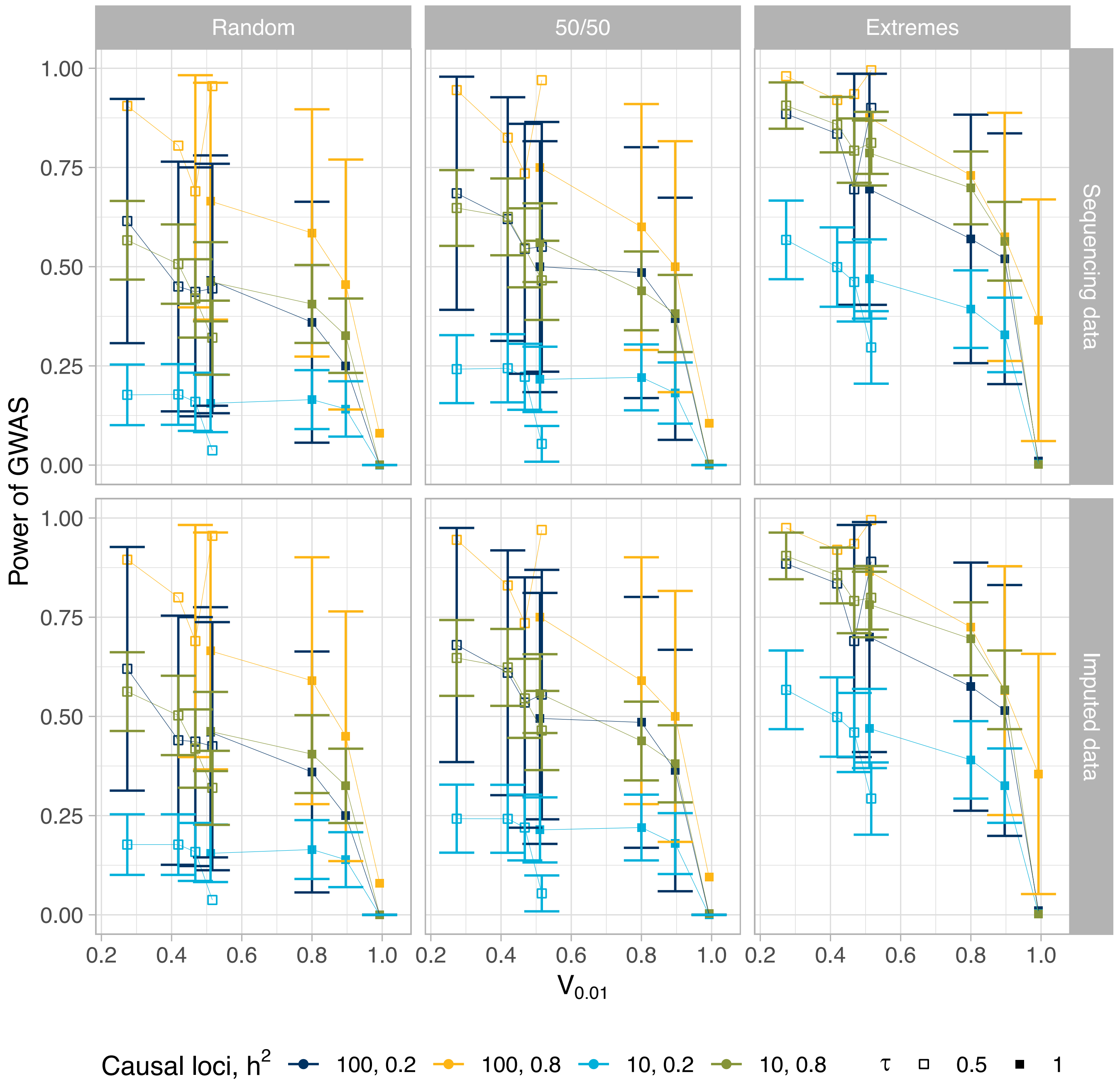


Figure S4. The statistical power of GWAS across different sampling strategies (columns) and across different sequencing methods (rows), as a function of the proportion of genetic variance explained by that genetic architecture at MAF=1%. Each point represents 20 independent simulations of 10 or 100 causal loci of 10kb each across a 5Mb simulated region for a given genetic architecture for a 50,000 individual African population.
